# All-optical electrophysiology reveals brain-state dependent changes in hippocampal subthreshold dynamics and excitability

**DOI:** 10.1101/281618

**Authors:** Yoav Adam, Jeong J. Kim, Shan Lou, Yongxin Zhao, Daan Brinks, Hao Wu, Mohammed A. Mostajo-Radji, Simon Kheifets, Vicente Parot, Selmaan Chettih, Katherine J. Williams, Samouil L. Farhi, Linda Madisen, Christopher D. Harvey, Hongkui Zeng, Paola Arlotta, Robert E. Campbell, Adam E. Cohen

**Affiliations:** Dept. of Chemistry and Chemical Biology, Harvard University; Dept. of Chemistry, University of Alberta; Dept. of Stem Cell and Regenerative Biology, Harvard University; Dept. of Neurobiology, Harvard Medical School; Allen Institute for Brain Science; Howard Hughes Medical Institute

## Abstract

A technology to record membrane potential from multiple neurons, simultaneously, in behaving animals will have a transformative impact on neuroscience research^1^. Parallel recordings could reveal the subthreshold potentials and intercellular correlations that underlie network behavior^2^. Paired stimulation and recording can further reveal the input-output properties of individual cells or networks in the context of different brain states^3^. Genetically encoded voltage indicators are a promising tool for these purposes, but were so far limited to single-cell recordings with marginal signal to noise ratio (SNR) *in vivo*^4-6^. We developed improved near infrared voltage indicators, high speed microscopes and targeted gene expression schemes which enabled recordings of supra- and subthreshold voltage dynamics from multiple neurons simultaneously in mouse hippocampus, *in vivo*. The reporters revealed sub-cellular details of back-propagating action potentials, correlations in sub-threshold voltage between multiple cells, and changes in dynamics associated with transitions from resting to locomotion. In combination with optogenetic stimulation, the reporters revealed brain state-dependent changes in neuronal excitability, reflecting the interplay of excitatory and inhibitory synaptic inputs. These tools open the possibility for detailed explorations of network dynamics in the context of behavior.

## Main text

Archaerhodopsin-derived genetically encoded voltage indicators (GEVIs) are excited by red light and emit in the near infrared part of the spectrum, have sub-millisecond response times at physiological temperature, and report neuronal action potentials with a ∼40% change in fluorescence brightness ^7-9^. A transgenic mouse expressing the Arch-based GEVI QuasAr2 and the channelrhodopsin CheRiff (together called ‘Optopatch2’) enabled optical stimulation and recording in sparsely expressing acute brain slices^4^, but did not attain adequate SNR for voltage imaging in brain *in vivo*. The primary factors which limited SNR were (a) poor trafficking of QuasAr2 *in vivo*, leading to intracellular aggregates that contributed to fluorescence background, and (b) high background, due to light scattering, out-of-focus fluorescence, and fluorescence from optically unresolvable neuropil.

To address the trafficking challenge, we systematically tested the impacts of trafficking sequences, appended fluorescent proteins, linker sequences, and various point mutations (*Methods*). We used the Optopatch system^7^ to test the SNR for optogenetically induced action potentials first in cultured neurons, and then in acute brain slices (Fig. S1). Incorporation of multiple K_ir_2.1 trafficking sequences, a Citrine fluorescent protein fusion (instead of mOrange2), and mutation of a putative ubiquitination site (lysine 171) to arginine, led to a construct that showed high expression and excellent trafficking *in vivo*, which we called QuasAr3 (Fig. 1). In acute cortical slices, QuasAr2 produced bright intracellular puncta, while QuasAr3, expressed under the same conditions, did not (Fig. 1a). Sparsely expressed QuasAr3 showed efficient trafficking throughout the dendritic arbor (Fig. 1b).

**Figure 1.**
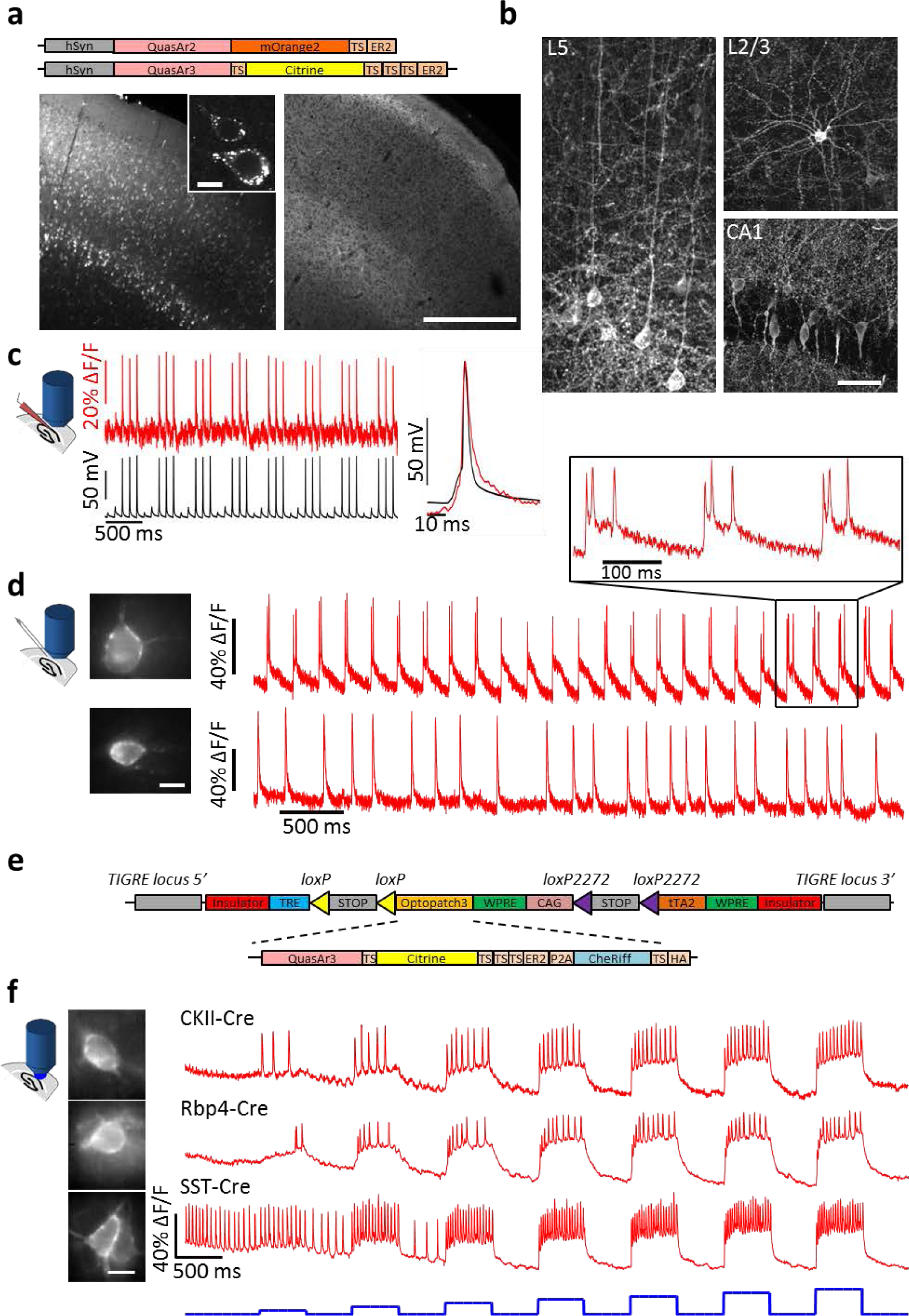
QuasAr3 shows improved membrane trafficking and reports action potentials in brain tissue. **(a)** Top: diagram of the QuasAr2 and QuasAr3 constructs. Bottom: Confocal images of brain slices expressing QuasAr2 and QuasAr3. Scale bar 500 μm. Inset: intracellular puncta in neurons expressing QuasAr2, scale bar 10 μm. **(b)** Confocal images of brain slices expressing Cre-dependent QuasAr3 with sparsity controlled by co-expression of hSyn-Cre. Scale bar 50 μm. **(c)** Simultaneous fluorescence and patch clamp recordings in acute brain slice show correspondence of optical and electrical signals. **(d)** Recordings from neurons expressing QuasAr3 using AAV virus in acute brain slice with activity evoked by field stimulation. **(e)** Construct design for a Cre-dependent Optopatch3 transgenic mouse. **(f)** All-optical electrophysiology recordings in acute brain slices from Optopatch3 transgenic mice crossed with different Cre driver lines. Detailed characterization of these mice is in Fig. S2. Scale bars in d, f 10 μm.

The protein engineering effort did not change the core amino acids relevant to voltage sensing. As expected, QuasAr3 retained the speed and sensitivity of QuasAr2 (Tables. S1 & S2). QuasAr3 reported action potentials in acute slice with close correspondence to patch clamp recordings and with high SNR (26.6 ± 4.8 in a 1 kHz bandwidth, n=5 neurons, Fig. 1c,d). We made a transgenic mouse with Cre-dependent QuasAr3 and CheRiff in the highly expressing TIGRE locus ^10^. Acute slices from these animals yielded high SNR genetically targeted all-optical electrophysiology recordings (Fig. 1e,f) and clearly showed cell type-specific differences in firing patterns (Fig. S2).

We then tested a point mutation in QuasAr3, V59A, whose homolog in bacteriorhodopsin (V49A) had previously been reported to cause photoswitching behavior ^11^ and to enhance the population of the fluorescent Q state^12^. QuasAr3(V59A) had an unusual optical property: under continuous red excitation (*λ*_exc_ = 640 nm, 10 W/mm^2^), illumination with moderate intensity blue light (*λ*_act_ = 488 nm, 100 mW/mm^2^) reversibly increased near infrared fluorescence (*λ*_em_ = 660 – 740 nm) by a factor of 2.9 ± 0.7, n = 8 cells (Fig. 2a). The enhancement did not occur in the absence of red light, nor in the parent QuasAr3 protein (Fig 2a, Fig. S3). Blue light also enhanced the amplitude of the voltage-dependent changes in fluorescence by a factor of 2.1 ± 0.5, n = 8 cells (Fig. 2 c). We called QuasAr3(V59A) ‘photoactivated QuasAr3’, or paQuasAr3.

**Figure 2.**
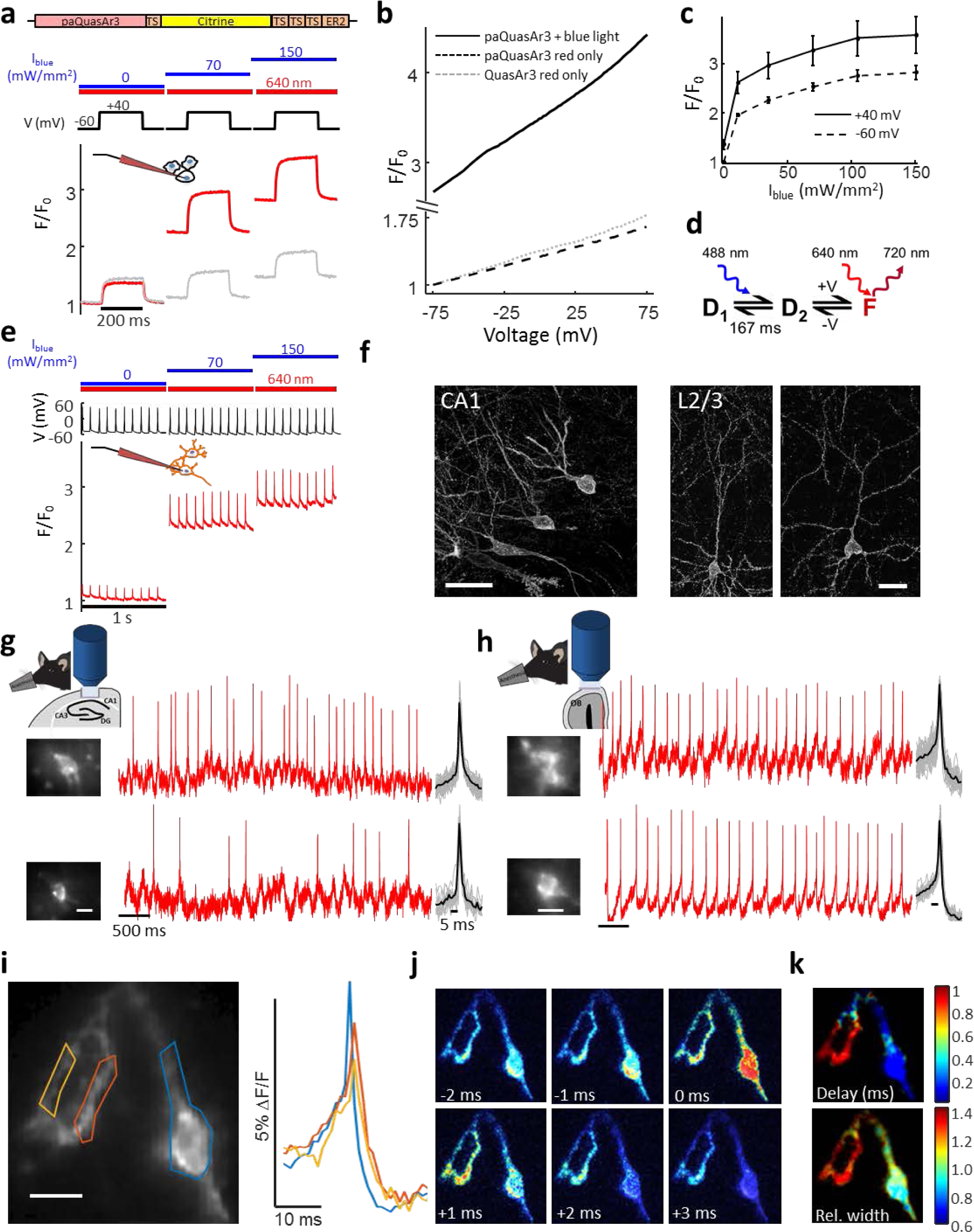
Photo-activated QuasAr3 (paQuasAr3) reports neuronal activity *in vivo*. **(a)** The V59A mutation in QuasAr3 rendered it photoactivatable by blue light. Red lines show near infrared fluorescence of a HEK cell expressing paQuasAr3 during voltage steps from -60 to +40 mV under constant red illumination (10 W/mm^2^) and variable blue illumination. Grey lines show the same experiment in a HEK cell expressing QuasAr3. **(b)** Voltage dependent change in near infrared fluorescence of paQuasAr3 and QuasAr3 with and without blue light (150 mW/mm^2^). **(c)** Dependence of paQuasAr3 photoactivation on blue light intensity at two membrane voltages. Error = mean ± s.e.m. **(d)** Model of the photocycle of paQuasAr3. Blue light converts the dark state, D_1_, into a pair of states, D_2_ and F, that show voltage-dependent near-infrared fluorescence. **(e)** Photoactivation enhanced voltage signals in a cultured neuron expressing paQuasAr3. **(f)** Maximum intensity projections of confocal stacks of sparsely expressed paQuasAr3 showed excellent trafficking in brain slices. **(g,h)** Optical measurements of paQuasAr3 fluorescence in (g) CA1 region of the hippocampus and (h) glomerular layer of the olfactory bulb of anesthetized mice. The red illumination (8 W/mm^2^, total power 10 mW) was structured to illuminate the soma of interest. Wide-field blue illumination was provided at 300 mW/mm^2^. **(i)** Spike-triggered average fluorescence from 88 spikes in a CA1 Oriens neuron. The fluorescence traces show clear delay and broadening of the spike waveform in the dendrites. **(j)** Frames from the spike-triggered average movie showing the delay in the back-propagating action potential in the dendrites relative to the soma. **(k)** Sub-Nyquist fitting of the action potential delay and width show electrical compartmentalization in the dendrites.

We characterized the kinetic, spectroscopic, and voltage-dependent properties of paQuasAr3. The blue light enhancement arose with a 50 ± 14 ms time-constant, subsided with a 167 ± 26 ms time-constant (n = 9 cells, Fig. S3), and showed saturation behavior, with 50% maximum enhancement at *I*_488_ nm = 27 mW/mm^2^ (Fig. S3). The absolute brightness of fully photoactivated paQuasAr3 (measured using fluorescence of the appended Citrine fluorophore to normalize for expression level) was 2-fold brighter than QuasAr3 (Fig S3). The photoactivation spectrum peaked at 470 nm, while the blue-sensitized fluorescence excitation spectrum had a similar shape to that of the parent QuasAr3, peaking at 580 nm (Fig. S4). The voltage-dependent properties were the same as in QuasAr3 (0.7 ms response time at 34°C, 56% ΔF/F per 100 mV, Tables S1 & S2, Fig. S3). Together, these findings suggested the photocycle model shown in Fig. 2d, in which blue light transferred population from a dark state, *D*_*1*_, to a voltage-sensitive equilibrium between a second dark state, *D*_*2*_, and a fluorescent state, *F*.

Cultured neurons expressing paQuasAr3 reported action potentials as spikes in fluorescence, and showed a 2.8 ± 0.2-fold enhancement in fluorescent spike amplitude upon blue illumination (n = 10, Fig. 2e). PaQuasAr3 expressed *in vivo* by viral transduction trafficked extremely well in soma and dendrites, with no detectable intracellular fluorescence (Fig. 2f). The ability to enhance the fluorescence signals 2.8-fold with targeted blue illumination, without affecting background, enabled fluorescence measurements of electrical activity from sparse paQuasAr3-expressing neurons *in vivo*. Fig. 2g,h shows representative epifluorescence recordings from single neurons in the CA1 region of the hippocampus and in the glomerular layer of the olfactory bulb of anesthetized mice.

In cells whose dendrites happened to lie within the focal plane of the soma, paQuasAr3 enabled sub-cellular mapping of action potential propagation. Fig. 2i-k shows an example of a CA1 Oriens interneuron undergoing spontaneous activity in an anesthetized mouse. A spike-triggered average movie, composed from 88 spikes, clearly showed a conduction delay between the cell body and the nearby dendrites. We used a sub-Nyquist interpolation algorithm^7^ to estimate the spike latency and width at each pixel in the image. These fits, performed independently at each pixel, showed clear electrical compartmentalization of the distal dendrites, in which the mean action potential was delayed and wider than in the soma (Fig. 2i-k; *Methods*), as anticipated from dendritic patch clamp recordings ^13^.

While the excellent membrane trafficking of paQuasAr3 enabled dendritic recordings, dense expression in acute slices resulted in broad and diffuse fluorescence, attributable to the efficient trafficking of the reporter into the optically unresolvable neuropil. This background overwhelmed signals from cell bodies, preventing functional imaging in densely expressing samples. Fusion of paQuasAr3 with a trafficking motif from the soma-localized K_V_2.1 potassium channel^14^,^15^ led to largely soma-localized expression (Fig. 3b). Confocal images of acute slices clearly resolved expressing neuronal cell bodies (Fig. 3b). We called this construct paQuasAr3-s.

**Figure 3.**
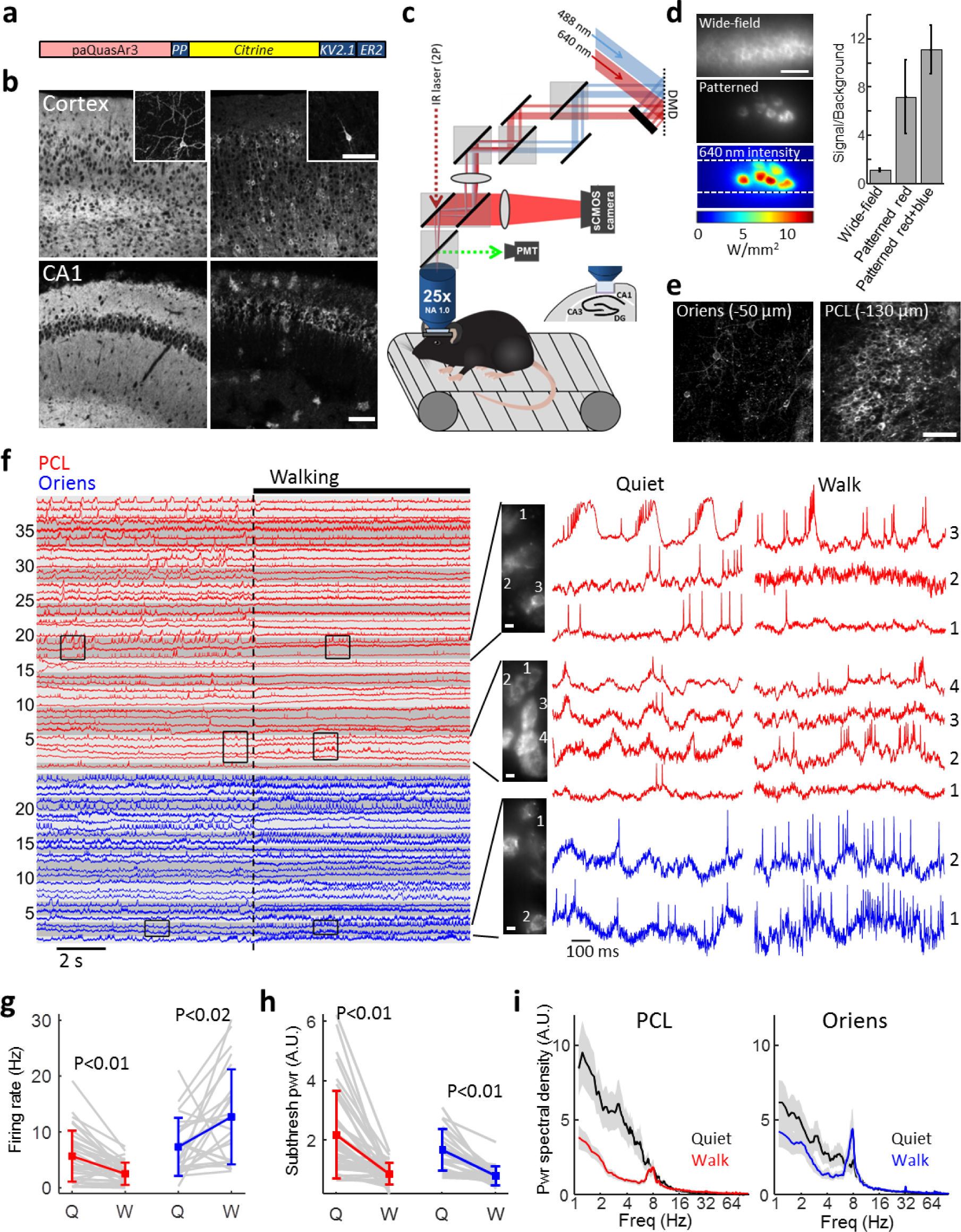
Optical recording of neuronal activity in hippocampus of walking mice. **(a)** Soma-localized paQuasAr3 construct (paQuasAr3-s). **(b)** Confocal images of brain slices show that highly expressed paQuasAr3 trafficked extensively in the neuropil, preventing optical resolution of individual cells (left), while paQuasAr3-s resolved cell bodies (right). Insets: sparsely expressed constructs showing the difference in dendritic expression between paQuasAr3 and paQuasAr3-s. Scale bars 100 μm. **(c)** Optical system for simultaneous 2-photon imaging, and patterned illumination with red and blue light. **(d)** Left, top: epifluorescence image with wide-field red illumination of densely expressed paQuasAr3-s in the CA1 region of the hippocampus, *in vivo*. Middle: epifluorescence image of the same field of view with patterned red illumination, showing clearly resolvable cell bodies. Bottom: pattern of red illumination and laser intensity. Scale bar 50 μm. Right: Effect of patterned red and blue light on signal-to-background ratio. **(e)** Two-photon images of paQuasAr3-s expression *in vivo*, in the Oriens (left) and the pyramidal cell layer (PCL, right). Scale bar 100 μm. **(f)** Fluorescence recordings from the PCL (red) and the Oriens (blue). Traces with similarly shaded backgrounds were acquired simultaneously. The traces show 9 s of quiet followed by 9 s of walking. Right: Magnified views of the traces showing complex spikes, bursts, correlated activity between cells, and modulation of the spiking activity by subthreshold dynamics. **(g)** Effect of brain state (Quiet or Walking) on firing rate in the PCL (red) and the Oriens (blue). **(h)** Effect of brain state on total power in the subthreshold oscillations in the PCL (red) and the Oriens (blue). While walking increased the mean firing rate in the Oriens, it decreased the power in the subthreshold oscillations. **(i)** Effect of brain state on population-average power spectra in the PCL and Oriens. Locomotion concentrated power in the θ-frequency band in both brain regions. Shading shows mean ± s.e.m.

Despite the soma-localized expression, wide-field epifluorescence images of densely expressing brain tissue showed a largely unstructured haze, a consequence of light scattering and out-of-focus fluorescence (Fig. 3d). To enhance the signal-to-background ratio *in vivo*, we developed a custom dual-wavelength micromirror-based illumination system to deliver both the red excitation and the blue sensitization light (Fig. 3c). Two-photon images of the appended Citrine fluorescent protein identified expressing cells. The red illumination was then patterned to impinge just on the cell bodies, avoiding interstitial regions. Patterned red illumination enhanced the signal-to-background ratio by a factor of 7.2 ± 2.3 (*n* = 10 cells, Fig. 3d). Red laser power into the tissue was 5-10 mW per cell. Adding blue illumination, also patterned to impinge only on the cell bodies (5-10 µW per cell), further enhanced the signal level by a factor of 1.6 (Fig. 3d).

We then made optical recordings from the hippocampus. Superficial cortex was removed ^16^ and recordings targeted either the Oriens layer (20 to 60 μm below the hippocampal surface) which comprises a sparse population of inhibitory neurons, or the pyramidal cell layer (PCL, up to 130 μm below the hippocampal surface) which comprises densely packed neurons, predominantly excitatory (Fig. 3e)^17^. In an anesthetized animal we recorded from up to 7 spiking cells simultaneously (Fig. S5). Optical recordings lasted up to 5 min with ∼50% photobleaching over this interval (Fig. S6).

To probe brain state-dependent changes in activity, we recorded from awake, head-fixed mice walking on a motorized treadmill, during 10 - 15 s of rest, followed by 15 s of walking (Methods). Recordings were made from *n* = 4 mice and probed 41 spiking neurons in the PCL from 14 fields of view and 24 spiking neurons in the Oriens from 14 fields of view (Fig. 3f). We used fluorescence transients associated with action potentials to segment the images into individual cells (Methods). Typical recordings resolved 2 – 4 spiking cells simultaneously in the PCL and up to 3 spiking cells simultaneously in the Oriens (Fig. 3f). All fluorescence traces are displayed at the native 1 kHz recording bandwidth, without smoothing.

Fluorescence traces clearly resolved hallmarks of CA1 intracellular activity, including simple and complex spikes, bursts, sub-threshold depolarizations, and θ-frequency oscillations (Fig. 3e)^18-21^. Visual inspection of the traces showed that spikes tended to ride atop subthreshold depolarizations, that subthreshold voltages were often correlated between nearby cells, and that many cells showed dramatic differences in subthreshold dynamics and spiking patterns between rest and walking. Fields of view containing multiple neurons could be re-identified and recorded in sessions up to 7 days apart (Fig. S7), opening the possibility to study long-term changes in subthreshold dynamics and intercellular correlations.

These recordings represent the first multicellular measurements of voltage dynamics in hippocampus of behaving mice. The data contain rich information on intra-and intercellular correlations and the effect of brain state (quiet vs. walking) thereon. We began the analysis by calculating basic statistics of the single-cell recordings. Forced walking decreased the mean spike rate of PCL neurons (5.7 ± 4.6 Hz to 2.5 ± 2.0 Hz, *p* = 9.7x10^-6^, *n* = 41 neurons), but, consistent with prior reports 22, increased the mean spike rate of Oriens neurons (7.3 ± 5.2 Hz to 12.7 ± 8.5 Hz, *p* = 0.01, *n* = 24 neurons, Fig. 3g). Walking decreased the overall power of the subthreshold oscillations in both the PCL (3.4 ± 4.3-fold reduction, *p* = 3 x 10^-6^) and Oriens (2.3 ± 1.5-fold reduction, *p* = 4 x 10^-6^, Fig. 3h), but increased θ-frequency power at 8 Hz in both layers (Fig. 3i), consistent with prior LFP recordings in freely walking animals^23^. Walking significantly decreased the rate of complex spikes (CS) in the PCL (0.3 ± 0.4 Hz to 0.04 ± 0.13 Hz, *p* = 5x10^-5^). As expected, CS were too rare to quantify in the Oriens interneurons (only one putative CS from all recordings)^19^.

Intercellular correlations in subthreshold voltage reflect shared synaptic inputs, an important measure of network architecture that is often not apparent from spike timing alone^24^. Paired patch recordings have quantified these correlations in cortex of behaving mice^25^,^26^, but not yet in the hippocampus. We therefore studied the pairwise cross-correlations in subthreshold voltage as a function of behavioral state (quiet vs. walking) and hippocampal layer (PCL vs. Oriens). Fig. 4 shows the auto- and cross-correlations of the fluorescence in trios of simultaneously recorded cells in both behavioral states and in both hippocampal layers. In the PCL the population-average equal-time cross-correlations were stronger when the mice were resting than when walking (0.29 ± 0.29 vs. 0.24 ± 0.26, mean ± s.d., *p* = 0.013, *n* = 40 pairs), while in the Oriens layer the mean strength of these correlations did not change (0.23 ± 0.33 vs. 0.23 ± 0.21, mean ± s.d., *p* = 0.99, *n* = 13 pairs). Similar behavior-dependent decreases in cross-correlation have been observed in excitatory cortical neurons ^25^, though the mean cross-correlations were lower in CA1 than previously reported in cortex. The degree of equal-time cross-correlation varied considerably between pairs of cells in both brain states (Fig. 4c,f). Most cell pairs showed positively correlated voltage fluctuations, though a small sub-population in each layer showed anti-correlated behavior (Fig. 4c,f, Fig. S8a).

**Figure 4.**
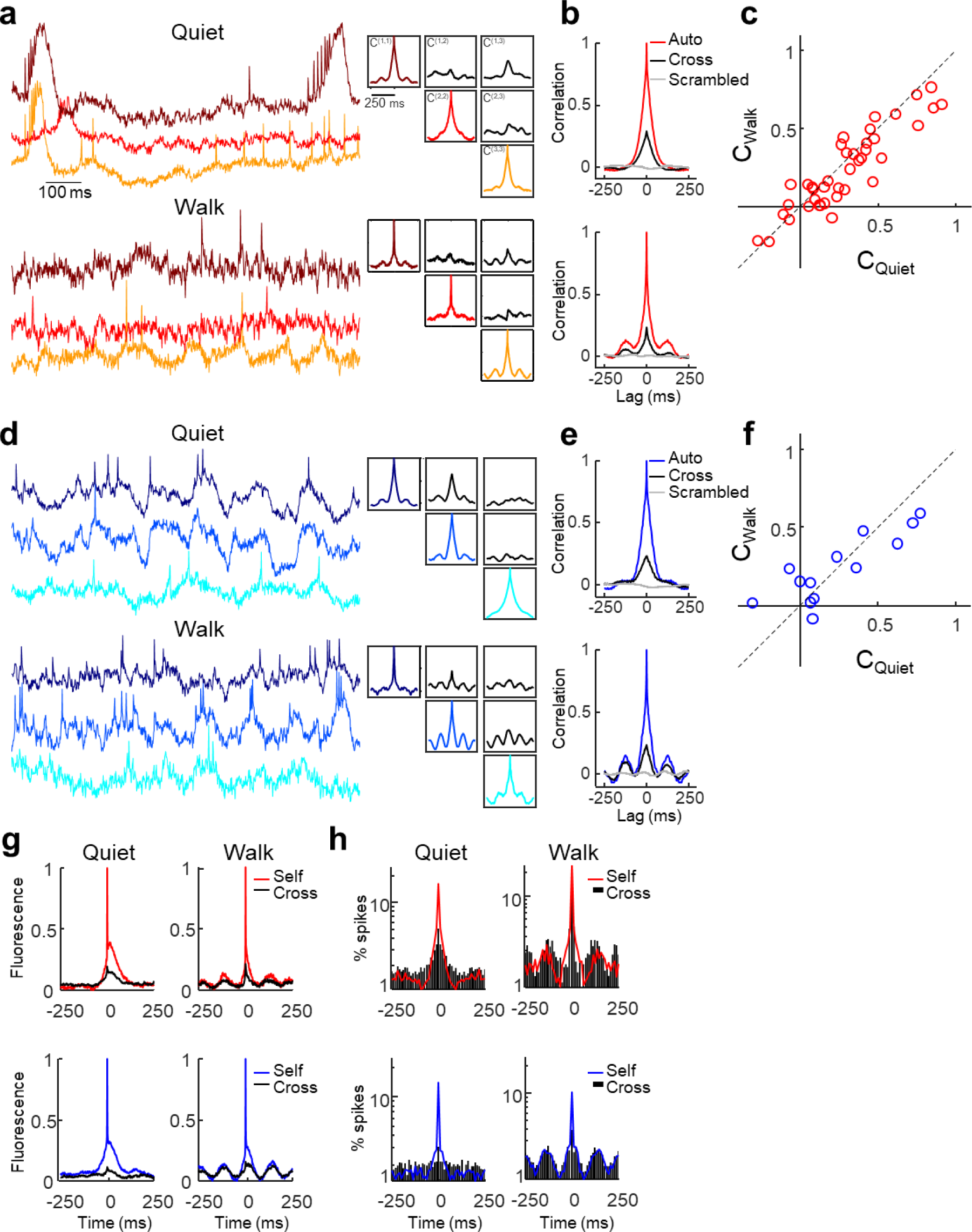
Behavior-dependent intercellular correlations in the hippocampus. **(a,d)** Left: Magnified sections showing 1 s recordings from a trio of cells in (a) the PCL and (d) Oriens. Top: recordings during quiet. Bottom: recordings from the same cells during walking. Right: auto- and cross-correlations of the fluorescence traces, calculated from 9 s recordings in each brain region and behavioral state. The auto-and cross-correlations clearly show enhanced θ-rhythm in both brain regions during walking and differing cross-correlations between simultaneously recorded pairs of cells. **(b,e)** Grand average auto- and cross-correlations. Mean cross-correlation between randomly selected cells from different fields of view shown in grey. **(c,f)** Distribution of equal-time correlation coefficients between pairs of simultaneously recorded cells. In the PCL correlations were significantly stronger during quiet than walking, as seen from the deviation from the dotted line. **(g)** Grand average spike-triggered average fluorescence during quiet (left) and walking (right) in the PCL (top) and Oriens (bottom). **(h)** Grand average spike-triggered average spiking probability during quiet (left) and walking (right) in the PCL (top) and Oriens (bottom). Y axis is on a log scale.

Little is known about the statistical structure of subthreshold voltages *in vivo*. Paired patch recordings *in vivo* in the cortex, have previously been decomposed into a shared subthreshold component, strongly correlated between the two cells, and a residuum, assumed to be independent between the cells ^1^,^25-28^. When more than two cells are measured simultaneously, an additional possibility arises: there could be multiple subthreshold signals, shared to varying degrees among different subsets of cells^29^,^30^. Dual recordings permit only one pairwise comparison, and thus cannot test this hypothesis. One must record subthreshold dynamics from three or more cells simultaneously, but to our knowledge no such recordings have been reported in any brain region *in vivo*.

If there were only one shared sub-threshold signal, the pairwise cross-correlation functions would have the same shape for all simultaneously recorded pairs of cells. Fig. 4 shows that in simultaneously recorded trios, the three pairwise cross-correlations can be distinctly different in shape and amplitude. In 9 FOVs in the PCL containing more than 2 cells, the cell-to-cell cross-correlation functions were only correlated between different pairs by a mean of 0.43 ± 0.57 during rest and 0.34 ± 0.51 during walking (*n* = 63 pairs of pairs), establishing that there exist multiple sub-threshold signals, shared to varying degrees among the cells. A challenge for future multi-cellular voltage imaging experiments will be to identify the underlying independent components and to assign their sources.

Spike-triggered averages (STA) of the fluorescence traces (Fig. 4g) reflect the relation between the sub-threshold inputs and the spiking output. In both the PCL and the Oriens, when a cell spiked, its STA membrane voltage (self-STA) had a positive-going deviation that preceded the spike. During walking, spikes in both layers occurred on average on the rising edge of the STA θ-rhythm, leading to a 35 degree phase shift in both layers between the mean spike and the peak of the mean intracellular θ- rhythm. We then considered the STA fluorescence of neighboring cells (cross-STA)^25-27^. Cross-STA waveforms were highly heterogeneous in both layers. Some pairs of cells showed nearly identical subthreshold components of self- and cross-STA, while others showed anti-correlated self- and cross- STA (see examples in Fig. S8). The grand average cross-STA waveforms showed a strong θ-rhythm component during walking, establishing that the shared θ-rhythm inputs strongly drove spiking.

Correlated sub-threshold potentials might lead to correlated spiking, though spike rates are often less correlated than are subthreshold voltages.^24^ STA plots of the spike probability (Fig. 4h) revealed that in the PCL, when a cell spiked, its neighbors also had a slightly elevated spike probability, likely a consequence of the shared subthreshold inputs. Strikingly, in the 150 ms surrounding a spike, the spiking cell tended to have a period of suppressed spike probability relative to its neighbors, consistent with prior reports^31^. Since the subthreshold voltages (Fig. 4g), reflecting synaptic inputs, did not show a relative hyperpolarization during this period, we ascribe the suppressed excitability around a spike to cell-autonomous decreases in excitability. In the Oriens, cells spiked at a relatively constant rate during the quiet period and at a rate modulated by the θ-rhythm during walking. In contrast to the PCL, when a cell spiked in the Oriens, neither it nor its neighbors showed perturbation to their spike rates at nearby times. These findings suggest that in the Oriens, spike rate was largely governed by an instantaneous rate process, strongly modulated by the shared subthreshold inputs.

Finally, we combined voltage imaging with simultaneous optogenetic stimulation to probe brain state-dependent changes in neuronal excitability in Oriens neurons. Voltage imaging alone cannot distinguish the relative contributions of excitatory and inhibitory inputs to the membrane potential. Paired optogenetic stimulation and voltage imaging open the possibility to probe changes in membrane resistance, reflecting shunting from inhibitory inputs.

We co-expressed soma-localized paQuasAr3 and soma-localized CheRiff from a single bicistronic vector (Fig. 5a) and applied steps of blue light illumination patterned only to the cell bodies (500 ms on, 500 ms off) at successively greater intensities (nominally 0.2 - 10 mW/mm^2^, not accounting for light scatter in tissue). Blue light-photoactivation of paQuasAr3 led to deterministic shifts in fluorescence baseline during periods of optogenetic stimulation which were readily corrected in post-processing (Fig. S9).

**Figure 5.**
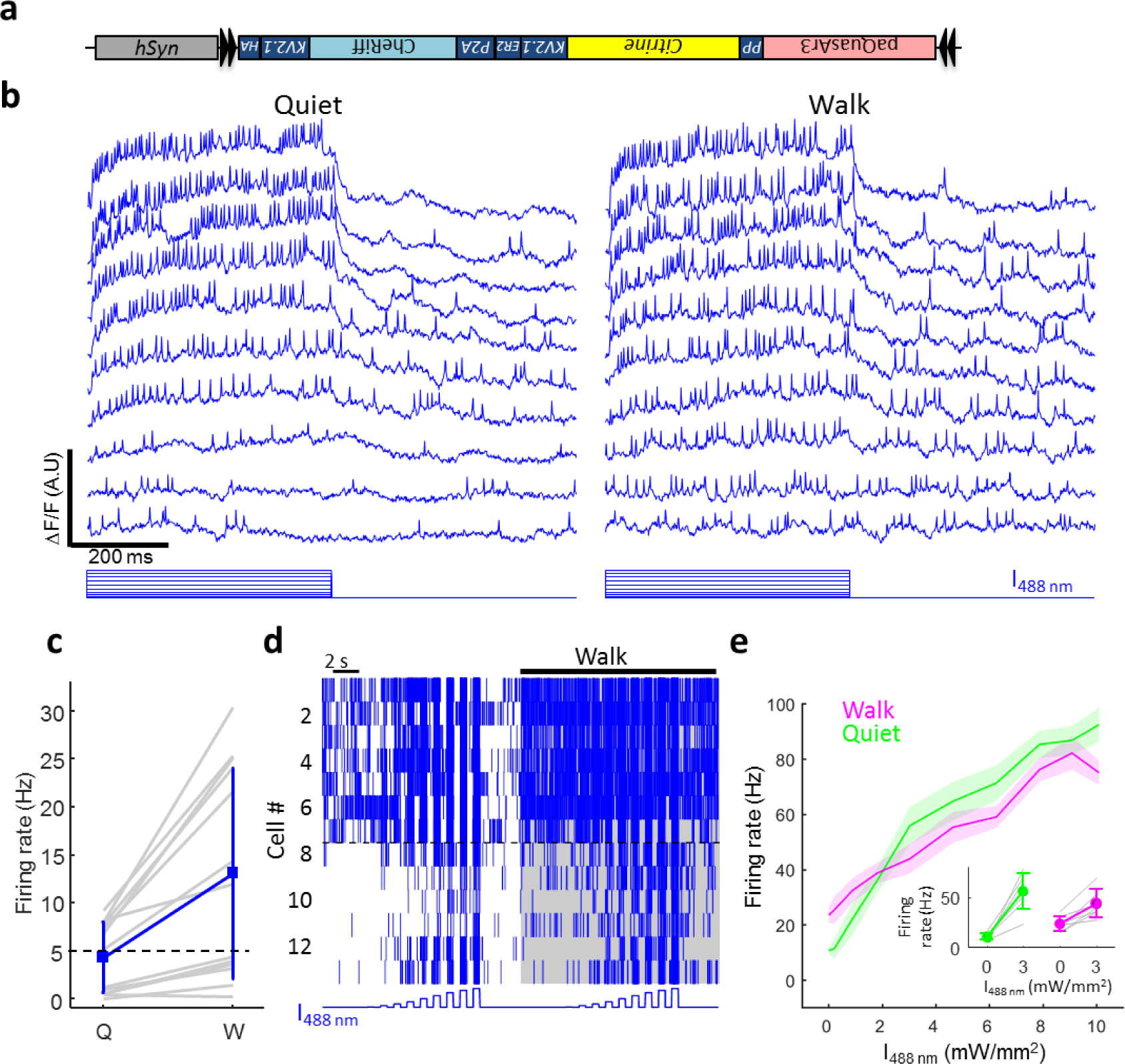
Simultaneous optogenetic stimulation and voltage imaging in hippocampus of walking mice. **(a)** Optopatch construct for viral Cre-dependent co-expression of CheRiff-s and paQuasAr3s. **(b)** Optically recorded activity of a single Oriens interneuron during optogenetic stimuli of different strength varying from 0 to 10 mW/mm^2^. Left: Quiet. Right: Walking. **(c)** Changes in firing rates between quiet and walking in absence of optogenetic stimulation. Gray: single cells. Blue: mean ± s.d. **(d)** Spike raster summarizing recordings from *n* = 13 interneurons. **(e)** Spike rate as a function of optogenetic stimulus strength during quiet and walking states. Analysis was restricted to the *n* = 7 cells that fired at > 5 Hz during quiet state, without optogenetic stimulation (dashed line in c & d). Shading shows mean ± s.e.m. Inset: increase in firing rate due to 3 mW/mm^2^ optogenetic stimulation in quiet and walking states. Gray: single cells. Colors: mean ± s.d.

Single cells showed enhanced firing under optogenetic stimulation (Fig. 5a). Cells typically showed erratic burst-like firing patterns, with spike rates up to 110 Hz. We then compared the relation of mean spike rate, *F*, to optogenetic stimulus strength, *I* (F-I curve) in the same cells during either quiet or walking behaviors. If optogenetic stimulation and synaptic inputs contributed additively to the overall spike rate, then the two F-I curves would be shifted along the x-axis but would have the same shape. Alternatively, if changes in inhibitory inputs led to changes in membrane resistance, then the slope of the F-I curves at weak stimulus would differ between the two conditions. Fig. 5e shows that the weak-stimulus slope of the F-I curve during walking was only 51 ± 27% of the slope during quiet (*n* = 7 cells, *p* = 0.007). The change in proportionality between optogenetic stimulus strength and firing rate implies a 2-fold decrease in membrane resistance during walking. These results show that changes in inhibitory inputs are an important contributor to the changes in activity during walking and demonstrate an all-optical technique for measuring excitation/inhibition balance *in vivo*.

Genetically encoded voltage indicators promise to be a powerful tool for studying neuronal dynamics *in vivo* in the context of behavior, but so far have been used mostly for technical demonstrations. To create a robust tool required improvements in signal-to-noise ratio in scattering brain tissue, which came from (a) improved membrane localization, (b) photo-activation, (c) soma localization, and (d) use of an optical system which localized illumination to the signal-generating regions of the sample. Despite these advances, robust voltage imaging was still restricted to the top ∼150 μm of the tissue. Improvements in GEVI brightness and photoactivation contrast will likely be needed to achieve greater depth penetration, and use of adaptive optical techniques may help combat tissue-induced aberrations.

The experimental system presented here opens many possibilities for exploring neural dynamics. By pairing voltage imaging with optogenetic activation or inhibition of distinct neural populations, one could probe the functional connectivity of different sub-populations, and probe the role of these populations in information processing. The ability to probe subthreshold dynamics over multiple days opens the possibility to study changes in circuit dynamics and intercellular correlations associated with learning and memory. Finally, simultaneous recordings of subthreshold voltages in large numbers of cells will reveal the underlying correlational structure which reflects the network architecture.

## Acknowledgments

We thank M.S. Lee and V. Joshi for help with tissue culture, D. Hochbaum, C. Straub, J.L. Saulnier, B.L. Sabatini, V. Kapoor and V. Murthy for help at early stages of the project. We thank A.H. Gheorghe for help with spectroscopy experiments, N. Rollins and S. Brownsberger for technical help and L. Yapp for advice on hippocampal surgeries. We thank A. Ruangkittisakul and K. Ballanyi for providing mouse hippocampus tissues for primary neuronal culture for the hierarchal screen. We thank G. Buzsaki and members of the Cohen lab for useful discussions. Y.A. was supported by a long term fellowship from the Human Frontiers Science Program and by a postdoctoral fellowship from the Edmund and Lili Safra center for Brain Sciences. This work was supported by the Howard Hughes Medical Institute.

## Author Contributions

Y.A. performed surgeries, *in vivo* imaging experiments, patch clamp and imaging experiments in acute slices and cultured neurons. J.J.K. performed patch clamp in HEK cells and developed the instrument control software. Y.A. did the protein engineering to optimize expression, membrane trafficking and soma localization. YZ, under the supervision of REC, did a parallel protein engineering effort with a hierarchical screen to test fusion proteins, linkers, and point mutations, which identified the beneficial trafficking effects of the V59A mutation. YA characterized mutants spectroscopically and electrophysiologically in mammalian cells. Y.A. designed and built the upright imaging system with help from S.K. and V.P. H.W. built the blue dual illumination path on the DMD. S.L. developed and characterized the Ai155 mice in collaboration with L.M. and H.Z. M.M.R. performed IUE, supervised by P.A. D.B. performed spectroscopy experiments in *E. coli*. S.C. and C.D.H. shared unpublished reagents related to soma targeting of opsins. K.J.W. helped with molecular biology. S.L.F. designed the CheRiff-HA construct. Y.A. and A.E.C. designed the project, analyzed data and wrote the manuscript. A.E.C. supervised all aspects of the project.

## Methods

## Protein engineering

### Screening pipeline

Performance of a genetically encoded voltage indicator (GEVI) depends on several independent parameters: brightness, sensitivity, speed of response, expression level and membrane trafficking. QuasAr2 showed excellent sensitivity and speed1, but dim fluorescence and poor membrane trafficking *in vivo* limited application *in vivo* (Fig. 1a). Trafficking efficiency and expression level can differ dramatically between cell types and growth conditions, so it is important to test candidate GEVIs in neurons.

We used the signal to noise ratio (SNR) for single spikes in cultured neurons as the screening parameter. The SNR integrates all relevant molecular features, and is the parameter most relevant to applications *in vivo*. To facilitate rapid evaluation of constructs we used the Optopatch configuration for simultaneous optical stimulation and optical measurement of action potentials, comprising bicistronic expression of the candidate GEVI with the blue shifted channelrhodopsin, CheRiff, using a P2A peptide^1^.

The starting material comprised QuasAr2 fused to dark mOrange2 (QuasAr2- dmOrange2-TS-ER2), driven by the CamKII promoter. We previously found that the presence of an appended fluorescent fusion was beneficial for trafficking^1^, but we wished to reserve the spectrum for other uses so we introduced the mutation Y71A into the mOrange2 backbone to destroy the chromophore^2^,^3^. TS represents the trafficking signal from K_ir_2.1^4^ and ER2 represents the endoplasmic reticulum export signal FCYENEV.^4^

QuasAr variants were then transfected into primary neurons using the calcium phosphate transfection method and tested for spike SNR using blue light stimulation (Fig. S1). Membrane trafficking *in vivo* was often worse than *in vitro* so constructs that showed SNR improvements in cultured primary neurons were then expressed *in vivo* using *in utero* electroporation (IUE), and tested for spike SNR in acute brain slices (Fig. S1).

### QuasAr3

The primary goal in developing QuasAr3 was to solve the membrane trafficking problem (Fig. 1a). We tested multiple modifications. Some other microbial rhodopsins (e.g. CheRiff, ChR86 and ChR1^5^) showed good membrane trafficking in rodent brain *in vivo*, so we explored chimeras comprising N-terminal fusions derived from these microbial rhodopsins. The N-terminus was added either appended to the QuasAr N-terminus or added after truncation of the first 16 residues from the QuasAr N-terminus. We also tried the N-terminal LUCY and LUCY-Rho tags which were shown to improve the surface expression of olfactory receptor^6^ and multiple repeats of the TS sequence^4^. Most modifications had neutral or negative effects on the SNR. Adding a second TS sequence in the linker between QuasAr2 and mOrange improved the spike SNR *in vitro* by 28%. Concatenating 3 TS sites instead of one at the C-terminus further improved SNR *in vitro* by 17% (Fig. S1d-f). Mutating a putative ubiquitination site in an intracellular loop (K171R) resulted in higher expression levels (quantified by dividing the0020QuasAr fluorescence with the fluorescence of the GFP fused to the co-expressed CheRiff, Fig. S1g).

In parallel to rational designs, we used a hierarchical screening approach for rapid evaluation of membrane trafficking and brightness of QuasAr point mutant and linker libraries (Fig. S1c). We found that lentiviral gene delivery was superior to conventional lipid-based transfection in primary cultured neurons as measured by homogeneity of expression, neuronal morphology and homogeneity of membrane trafficking. Therefore, we sought to develop a screening strategy in which constructs that showed good brightness in *E. coli* could immediately be packaged in lentiviral vectors and tested in neurons, without laborious re-cloning. We inserted a putative prokaryotic promoter sequence BBaJ23101: TTTACAGCTAGCTCAGTCCTAGGTATTATGC and an *E. coli* ribosome binding site (AAAGAAGAGAAA) between the CAMKII promoter and the Kozak sequence of plasmid FCK-Arch-GFP (Addgene #22217).^7^ The resulting vector is termed FCK_DuEx1.0 for gene expression in both prokaryotic and eukaryotic cells as well as lentivirus production.

We created libraries of QuasAr2 fused to either eGFP, Citrine, mKate, or mRuby with a linker containing 2 randomized amino acids between QuasAr2 and the FP fusion. For library screening, *E. coli* colonies were transformed with linker libraries in FCK_DuEx1.0. The colonies with the brightest fluorescence were picked for lentivirus production in HEK cells and secondary screening in primary neuronal culture. For lentivirus production, HEK cells were co-transfected with the QuasAr variants in FCK_DuEx1.0, along with packaging plasmids pCMV-dR8.2 dvpr (Addgene #8455) and pCMV-VSV-G (Addgene #8454). The media from the HEK cell culture was collected 48 hours after transfection and 10% of the lentivirus-containing media was added into the primary hippocampal neuronal cultures for transduction. After 2-5 days, the membrane trafficking of QuasAr mutants was evaluated using fluorescence microscopy. The best constructs in this screen included Citrine as the fluorescent protein fusion with 4 different linkers – GE, GP, PP or PP. These constructs were then cloned in the Optopatch configuration and tested for spike SNR as described above. Two of these linkers - PP and PT showed better SNR compared with QuasAr2-mOrange (Fig. S1e).

We next combined the best modifications into a new construct: QuasAr2(K171R)-TS-Citrine-TS-TS-TS-ER2 which we call QuasAr3. We packaged QuasAr3 into AAV and validated its performance in acute slices (Fig. 1).

### paQuasAr3

We explored the effects of previously described Arch point mutations T27A, V59A, T99C, P196S, I129V, I129T, and A225M^8^ on membrane trafficking, brightness and sensitivity of QuasArs 1 and 2 using the hierarchical screening pipeline described above. We introduced these mutations separately into QuasAr2-Citrine, and estimated brightness by expressing these constructs in primary neurons and dividing the QuasAr fluorescence by the Citrine fluorescence. Initially it seemed that none of the mutations improved the QuasAr brightness. However, when we tested mutation V59A for spike SNR using blue light optogenetic stimulation, we found that fluorescence of the V59A mutant was potentiated by blue light. The photoactivation property of the V59A mutant may be related to the photoswitching property observed in the homologous bacteriorhodopsin mutant (V49A).^9^

To our surprise, we found that the V59A mutation led to near perfect membrane trafficking. This improved trafficking may be related to the enhanced thermal stability observed in the homologous bacteriorhodopsin mutant.^9^ The photoactivation property, combined with the excellent trafficking, motivated us to further characterize QuasAr3(V59A) (Fig.2, Fig.S3, Fig.S4, Tables S1 & S2).

### paQuasAr3-s and CheRiff-s

The trafficking sequence of the K^10^

V2.1 channel was previously shown to restrict the expression channelrhodopsin 2 to the soma and proximal dendrites ^10^,^11^. To produce soma-localized paQuasAr3 and CheRiff, we removed all the K_ir_2.1 trafficking sequence (TS) and introduced the K_V_2.1 sequence: QSQPILNTKE MAPQSKPPEE LEMSSMPSPV APLPARTEGV IDMRSMSSID SFISCATDFP EATRF either in the linker, at the C-terminus, or in both positions. In the QuasAr construct with a single K_V_2.1 sequence in the C-terminus, we used PP for the linker (see above). CheRiff was directly fused with GFP. These 6 constructs were then expressed in primary hippocampal neurons and visually inspected for localization. For both CheRiff and paQuasAr3, expression was most soma-restricted when the K_V_2.1 motif was at the C-terminus. These constructs were further characterized both in slices and *in vivo* (Fig3, Fig5).

### Spectroscopic studies

To probe the spectroscopic properties of the QuasAr3 and paQuasAr3 proteins, we expressed them in *E. coli* as described before ^1^. Briefly, *E. coli* cultures were grown overnight in liquid LB medium with 200 μg/mL ampicillin. The next day the cultures were supplemented with 50 µM all-trans-retinal and 0.1% arabinose and incubated for 4 hours at 37°C. Cell pellets were collected by centrifugation and kept in -80°C until use. We measured the excitation and photoactivation spectra using a previously described microscope equipped with a wavelength-tunable supercontinuum light source and an EMCCD detector ^12^.

## Optical systems

### Imaging HEK cells and primary neurons

Experiments were conducted on a home-built inverted fluorescence microscope equipped with 405 nm, 488 nm, 532 nm, and 640 nm laser lines and a scientific CMOS camera (Hamamatsu ORCA-Flash 4.0). Laser beams were combined using dichroic mirrors and sent through an acousto-optic tunable filter (AOTF; Gooch and Housego TF525-250-6-3-GH18A) for temporal modulation of intensity of each wavelength. The beams were then expanded and focused onto the back-focal plane of a 60× water immersion objective, numerical aperture 1.2 (Olympus UIS2 UPlanSApo 60×/1.20 W). Imaging of fluorescent proteins was performed at illumination intensities of 20-40 mW/mm^2^. Imaging of QuasAr fluorescence was performed at an illumination intensity of 4 W/mm^2^. Stimulation of CheRiff was performed at an illumination intensity of 0.4-1.2 mW/mm^2^. For fast data acquisition, a small field of view around the cell of interest was chosen at the center of the camera to achieve a frame rate of 1 kHz.

### Imaging acute slices and live mice

Experiments were conducted on a home-built upright fluorescence microscope equipped with 488 nm and 640 nm laser lines and a scientific CMOS camera (Hamamatsu ORCA-Flash 4.0). The 488 nm line was sent through an acousto-optic tunable filter (Gooch and Housego 48058-2.5-.55) for intensity modulation, and then expanded and focused onto the back-focal plane of the objective.

In the first generation of the microscope, we used 140 mW 640 nm laser expanded to a single ∼50 μm circular spot, and a 20x NA1.0 objective (Olympus XLUMPLFLN 20×/1.0 W). This configuration was used for the acute brain slice experiments (Fig. 1b,d) and olfactory bulb *in vivo* experiments (Fig. 2h). Data were acquired on a small field of view around the cell of interest at a frame rate of 1 kHz.

In the second generation of the microscope, we used an 8 W, 640 nm diode bar laser (DILAS MB-638.3-8C-T25-SS4.3) and patterned the illumination using a digital micromirror device (DMD, Vialux, V-7000 UV, #9515) to illuminate specific sub-cellular structures and to avoid illuminating interstitial regions of the brain slice. We used a 16x 0.8 N.A. objective with 3 mm working distance (Nikon CFI75 LWD 16xW). Sampling rate was 1 kHz. This configuration was used for the single cell *in vivo* measurements in the hippocampus (Fig. 2 g,i-k).

In the third generation of the microscope, we patterned both the 488 nm and 640 nm lasers to achieve overlapping red and blue spots in the sample. The beams were then combined on a dichroic beamsplitter and aligned to be side-by-side and propagating parallel. The combined beam was then sent to a DMD, so half of the DMD patterned one wavelength and half of the DMD patterned the other wavelength. The two wavelengths were then split and recombined with a pair of dichroic mirrors to form a single beam with independently patterned blue and red excitation (Fig.3c). Cylindrical lenses were used to correct for aberrations introduced by slight warping of some dichroic mirrors.

The patterned epi-illumination was combined with a home-built scanning two photon system. The visible and near-infrared beams were combined using an 875 nm long-pass dichroic mirror (Semrock). A 532 nm notch dichroic mirror (Semrock) directed green emission (from Citrine or GFP) to a photomultiplier, while near infrared fluorescence (from QuasAr) was sent to a Hamamatsu sCMOS camera. This configuration was used for all multicellular recordings in the hippocampus *in vivo* (Figures 3 - 5) using a 25x NA1.0 objective with 4 mm working distance (Olympus XLPLN25XSVMP2). Red laser intensity was 12 W/mm^2^ and sampling rate was 1 kHz. This configuration was also used for the measurements from brain slices of transgenic mice (Fig. 1f and Fig. S2). These data were acquired with a 20x objective (Olympus XLUMPLFLN 20×/1.0 W), red laser intensity 20 W/mm^2^ and sampling rate 500 Hz.

## Primary neuronal culture and gene delivery

All procedures involving animals were in accordance with the National Institutes of Health Guide for the care and use of laboratory animals and were approved by the Harvard Institutional Animal Care and Use Committee (IACUC).

Hippocampal neurons from P0 rat pups were dissected and cultured on rat glial monolayers as described before^1^. Briefly, P0 neurons were plated in neurobasal-based medium (NBActiv4, BrainBits LLC.) at a density of 30,000–40,000 cm^-2^ on the pre-established glial monolayers. At one day *in vitro* (DIV), cytarabine was added to the neuronal culture medium at a final concentration of 2 µM to inhibit further glial growth.

Neurons were transfected between DIV 7-10 via the calcium phosphate transfection method. Measurements on neurons were taken between DIV 14-18.

## Imaging and electrophysiology in HEK cells and primary neurons

HEK293T cells were cultured and transfected as described before.^1^ Briefly, HEK-293 cells were grown at 37 °C, 5% CO_2_, in DMEM supplemented with 10% FBS and penicillin-streptomycin. 200-400 ng of plasmid DNA was transfected using Transit 293T (Mirus) following the manufacturer’s instructions, and assayed 48 hours later. The day before recording, cells were re- plated onto Matrigel coated glass-bottom dishes (In Vitro Scientific) at a density of ∼10,000 cells/cm^2^.

All imaging and electrophysiology were performed in extracellular buffer containing (in mM): 125 NaCl, 2.5 KCl, 3 CaCl_2_, 1 MgCl_2_, 15 HEPES, 30 glucose (pH 7.3) and adjusted to 305-310 mOsm with sucrose. A gap junction blocker, 2-aminoethoxydiphenyl borate (50 µM, Sigma), was added to HEK cells to eliminate electrical coupling between cells. Primary neurons were supplemented with excitatory synaptic blockers (either 20 μM NBQX or 20 μM CNQX, both from Tocris).

Optopatch measurements and the simultaneous whole-cell patch clamp and fluorescence recordings were acquired on the home-built, inverted epifluorescence microscope described above. For simultaneous electrophysiology and imaging, filamented glass micropipettes (WPI) were pulled to a tip resistance of 5–10 MΩ, and filled with internal solution containing (in mM):125 potassium gluconate, 8 NaCl, 0.6 MgCl_2_, 0.1 CaCl_2_, 1 EGTA, 10 HEPES, 4 Mg-ATP, 0.4 Na-GTP (pH 7.3); adjusted to 295 mOsm with sucrose. Pipettes were positioned with a Sutter MP285 manipulator. Whole-cell, voltage and current clamp recordings were acquired using an Multiclamp 700B amplifier (Molecular Devices), filtered at 2 kHz with the internal Bessel filter and digitized with a National Instruments PCIE-6323 acquisition board at 10 kHz.

## Gene targeting in ES cells and generation of knock-in Cre-dependent reporter mice

To generate the Ai155 targeting vector, Optopatch3 comprising QuasAr2(K171R)-TS-Citrine- TSX3-ER2-P2A-CheRiff-TS-HA was inserted into the cre-dependent TIGRE 2.0 construct13. Gene targeting was performed at the Allen Institute for Brain Science. Targeting of the transgene cassettes into the TIGRE locus was accomplished via Flp-recombinase mediated cassette exchange (RMCE) using circularized targeting vector, a CAG-FlpE vector (Open Biosystems), and a Flp recombinase landing pad ES cell line derived from G4 cells^14^. Correctly targeted ES cells were identified via PCR, qPCR, and Southern blots. ES clones were karyotyped and verified to be chromosomally normal. ES clone injection was performed at the Harvard University Genetic Modification Facility. Optopatch-positive ES clones were injected into C57BL/6J blastocysts to obtain chimeric mice. Chimeric mice were bred with C57BL/6J mice to obtain F1 Floxopatch+/- mice. The Ai155 mouse line is on a mixed C57BL/6J; C57BL/6N genetic background. The Ai155 mouse line has been deposited in Jackson lab with Stock No: 029679.

Ai155 mice were crossed with the following Cre driver lines: CKII-Cre (gift from Venkatesh Murthy), Somatostatin (SST)-IRES-Cre (JAX #013044) and Rbp4-Cre (MMRRC- 031125-UCD, gift from Bernardo L. Sabatini,)

### Genotyping

The presence of QuasAr3 was determined with the primer pair: 5’- GCTGGTCTCCAACTCCTAATC -3’ 5’-CTGTATCTGGCTATGGCCG -3’. A 1.07 kb amplicon indicated presence of the gene. Homozygous mutant insertion was determined with the primer triplets: 5’- GTG TAG CCC TGG CTT TTC TG-3’, 5’-GAA CTC ACA GTG GCC AGT CA-3’, 5’- TCC CCT GGC ACA ACG TAA G-3’. These yielded a 295 bp amplicon for mutant band and a 468 bp amplicon for wild type band.

## Imaging and electrophysiology in acute slices

### In utero electroporation (IUE)

In utero electroporation was performed in timed pregnant CD-1 mice as previously described.^15^ The day of vaginal plug was designated as embryonic day 0.5 (E0.5), while the day of birth was designated as postnatal day 0 (P0). Briefly, 1 µL of endonuclease-free purified DNA (3.75 µg/µL comprising 2.75 µg/µL of the candidate GEVI construct and 1 µg/µL of tdTomato control) in sterile PBS mixed with 0.005% Fast Green was injected into the lateral ventricle of embryonic day 14.5 mice under ultrasound guidance (Vevo 770, VisualSonics). Five 35 volt pulses of 50 ms duration at 1 s intervals were delivered outside the uterus in appropriate orientation using 1 cm diameter platinum electrodes and a CUY21EDIT square-wave electroporator (Nepa Gene). Tested constructs were co-electroporated with either CBIG-tdTomato plasmid (gift from Jeffery Macklis) or pCAG-GFP (gift from Connie Cepko, also available as Addgene 11150) for visual identification of electroporated embryos using a fluorescence stereoscope.

### AAV virus preparation

Constructs were cloned into an AAV transfer plasmid either with hSyn promoter (Addgene #51697) or for Cre-dependent expression with the CAG promoter (Addgene #22222)^7^. In order to pack the Cre-dependent Optopatch constructs into the 4.5 kb size limitation of AAV, we had to modify the both the construct and the transfer plasmid. First, we replaced the GFP fusion of CheRiff with a short HA tag. Second, we used a Cre-dependent transfer plasmid with an hSyn promoter instead of CAG (Addgene #44362) ^16^. Third, we swapped the 480 bp bGH polyA sequence with 120bp SV-40 polyA.. See table S3 for details of the AAV vectors used in this study. All AAV plasmids were deposited with Addgene. Viruses were produced by the Gene Transfer Vector Core at Massachusetts Eye and Ear Infirmary & Schepens Eye Research Institute (MEEI), Harvard Medical School. All experiments used AAV serotype 2/9.

### Virus injection for acute slices measurement

For acute brain slice experiments (Fig. 1a-d), we injected AAV virus in C57BL/6 P0-P1 mice. Virus was diluted and injected at final titers of 1x10^13^ GC/mL. To achieve sparse expression, Cre dependent constructs were mixed with hSyn-Cre virus at final titers of 1 to 5x10^10^ GC/mL.

P0-P1 pups were cryo-anesthetized and immobilized dorsal side up under a stereotaxic setup. Injections were made using home-pulled micropipettes (Sutter P1000 pipette puller), mounted in a microinjection pump (World Precision Instruments Nanoliter 2010) controlled by a microsyringe pump controller (World Precision Instruments Micro4). The micropipette was positioned using a stereotaxic instrument (Stoelting). Pups were injected in the left hemisphere, 0.9 mm lateral and 0.9 mm anterior to lambda. Starting at a depth of 1.0 mm beneath the surface of the skull, virus injections (40 nL, 5 nL/s) were performed at 0.1 mm increments as the pipette was withdrawn. Pups were placed back in their home cage once they were awake.

### Acute slice preparation

Acute brain slices were prepared from P16–P28 mice as described before.^2^ Briefly, mice were anesthetized with isoflurane, and then subjected to intracardiac perfusion with ice-cold slicing solution containing (in mM): 110 choline chloride, 2.5 KCl, 1.25 NaH_2_PO_4_, 25 NaHCO_3_, 25 glucose, 0.5 CaCl_2_, 7 MgCl_2_, 11.6 Na-ascorbate, and 3.1 Na-pyruvate and saturated with carbogen (95% O_2_, 5% CO_2_). Mice were then decapitated and the brains were rapidly dissected and sliced into 300 μm coronal sections using a vibratome (Leica VT 1200S). Mice < P21 were directly decapitated without intracardiac perfusion. Slices were incubated for 45 min at 34 °C in a carbogenated artificial CSF (ACSF) containing (in mM): 127 NaCl, 2.5 KCl, 1.25 NaH_2_PO_4_, 25 NaHCO_3_, 25 glucose, 2 CaCl_2_, and 1 MgCl_2_. The osmolarity of all solutions was adjusted to 300–310 mOsm and the pH was maintained at 7.3 under constant bubbling with carbogen.

### Imaging and electrophysiology in acute slices

Acute slices were imaged on a home built upright microscope described above. Measurements were conducted in carbogen-saturated ACSF at room temperature. ACSF was perfused at a rate of 2 mL/minute. For simultaneous imaging and whole cell patch clamp measurements, filamented glass micropipettes (WPI) were pulled to a tip resistance of 4-6 MΩ, and filled with internal solution containing (in mM): 125 potassium gluconate, 8 NaCl, 0.6 MgCl2, 0.1 CaCl2, 1 EGTA, 10 HEPES, 4 Mg-ATP, 0.4 Na-GTP (pH 7.3); adjusted to 295 mOsm with sucrose. Pipettes were positioned with a Sutter MP285 manipulator. Whole-cell, current clamp recordings were acquired using an MultiClamp 700B amplifier (Molecular Devices), filtered at 2 kHz with the internal Bessel filter and digitized with a National Instruments PCIE-6323 acquisition board at 10 kHz

### Confocal imaging

Acute slices were fixed in 4% paraformaldehyde (PFA) and confocal fluorescence imaging was performed on an Olympus FV1000 confocal microscope at the Harvard Center for Brain Sciences microscope facility.

## Cranial windows, virus injections, training, and *in vivo* imaging

### Virus injection and cranial window surgery

8-12 week old C57BL/6 mice (male and female) were deeply anesthetized with 2% isoflurane and maintained with ∼1% isoflurane throughout the surgery. The skull was exposed and thoroughly dried and a 3 mm round craniotomy was opened using a biopsy punch (Miltex). For Olfactory bulb (OB) imaging, the craniotomy covered both OBs as previously described17. For CA1 imaging, the craniotomy center was 1.8 mm lateral, 2.0 mm caudal of bregma. Virus was then injected in 1-3 locations in the center of the craniotomy.

For single-cell measurements with paQuasAr3 (Fig. 2), sparse expression was achieved by mixing CAG-FLEX-paQuasAr3 AAV virus (final titer 2x10^13^ GC/mL) with hSyn-Cre virus (final titer 2x10^10^ to 1x10^11^ GC/mL). For dense expression of paQuasAr3-s, we used low titer CAG-FLEX-paQuasAr-s virus (final titer 6x10^11^ GC/mL) or hSyn-DiO-paQuasAr3-s-P2A- CheRiff-s (final titer 1.6x10^12^ GC/mL), mixed with higher titer CKII(0.4)-Cre virus (UPenn vector core, final titer 7x10^11^ GC/mL). Injections were made using home-pulled micropipettes (Sutter P1000 pipette puller), mounted in a microinjection pump (World Precision Instruments Nanoliter 2010) controlled by a microsyringe pump controller (World Precision Instruments Micro4). The micropipette was positioned using a stereotaxic instrument (Stoelting Digital Mouse Stereotaxic Instrument). For CA1 expression, injections were made -1.5 mm to -1 mm from Dura, at 0.1 mm increments (40 nL per depth, 5 nL/s). For OB expression, injections were made from -0.4 to 0 mm from dura at 0.1 mm increments (40 nL per depth, 5 nL/s). Brain surface was kept moist with saline throughout the injection.

In the OB, a 3 mm round #1 cover glass (Harvard apparatus) was then placed on the OB surface, and sealed with cyanoacrylate and dental cement (C&B Metabond). A titanium bar was glued behind the window, skin was sutured and animal was sent to recovery.

The procedure for imaging in CA1 followed Dombeck and coworkers.^18^. Briefly, a cannula was prepared prior to the surgery and comprised a 1.5 mm segment of a 3 mm outer diameter thin walled stainless steel tube (MicroGroup). A 3 mm diameter #1 round coverglass (Harvard apparatus) was cemented to one end of the tube using UV curable adhesive (Norland Products) and cured for at least 5 minutes on a standard lab UV table. Following hippocampal virus injection, we removed the dura, and then slowly aspirated the cortex while continuously irrigating with saline until bleeding stopped. After exposure of the external capsule, a small region of the capsule in the center was gently removed, exposing the CA1 surface. To reduce brain motion during locomotion, a small amount of Kwiksil (WPI) was applied to the surface of the brain and the cannula was then inserted and cemented to the skull with dental cement (C&B metabond). After the cannula cured, a titanium headplate (similar to Ref. 19) was glued around the cannula and any exposed skull was covered with dental cement. Animals were returned to their home cage for recovery and treated for 3 days with Carprofen (5 mg/kg) and Buprenorphine (0.1mg/kg) twice a day. To avoid damage to the implant, mice were housed in separate cages.

### Imaging anesthetized animals

Imaging typically started 3 weeks post-surgery. Mice were lightly anesthetized (0.7-1% isoflurane) and head-fixed under the upright microscope (see above) using the titanium head plate. Eyes were kept moist using ophthalmic eye ointment. Body temperature was continuously monitored and maintained at 37°C using a heating pad (WPI). A typical imaging session lasted 1-2 hours, and then animals quickly recovered and returned to their home cage.

### Motorized treadmill

A home built treadmill was composed of two 3D printed plastic wheels (5 cm wide, 10 cm in diameter). We used a 5 cm wide, 180 cm long velvet belt (McMaster Carr #88015K1). Treadmill speed was regulated using a computer controlled small electric motor (Pololu #3042). Linear speed was 5 to 10 cm/s.

### Imaging awake, walking animals

Head-fixed animals were imaged while walking on a home-built motorized treadmill. Before imaging sessions started, mice were habituated to head restraint and walking by training them at least 3 times, every 24 hours, for 15-30 minutes. During each training session mice were first habituated to head restraint until completely relaxed, and then started walking on the treadmill. Each walking period lasted 1 minute followed by at least 2 minutes of rest. Walking speed was 5 to 10 cm/s. To adjust mice to imaging conditions, at least one training session took place under the microscope with the objective on top of the cranial window.

For experimental runs, we first used a protocol of 10 s rest following by 3x 15 s periods of increased speed (5, 7.5 and 10 cm/s) followed by additional 10 s rest (65 s in total, Fig. S7). We observed that the difference in firing patterns was more starkly different between rest and walking than between different walking speeds, so we switched to a protocol of 15 s rest followed by 15 s walking at 10 cm/sec repeated twice (60 s in total). In each field of view (FOV), we repeated the protocol 2 to 5 times. Imaging session lasted up to 1 h and then animals were returned to their home cages.

## Data analysis

### Statistics

Statistical tests were performed using standard Matlab functions (Mathworks). All error bars represent standard deviation unless otherwise noted. For two-sample comparisons of a single variable, a two-sided Student’s *t*-test (paired or unpaired) was used.

### Extracting fluorescence from single cell movies

First, an estimate of the photobleaching baseline was constructed from the mean intensity of the whole FOV by applying sliding minimum filter, followed by a sliding mean filter. Each frame of the movie was then divided by this baseline. Fluorescence values were extracted from corrected movies in one of two ways. When single cells were well isolated from other fluorescent sources, we used the maximum likelihood pixel weighting algorithm described in Kralj *et al.*^20^ Briefly, the fluorescence at each pixel was correlated with the whole-field average fluorescence. Pixels that showed stronger correlation to the mean were preferentially weighted. This algorithm automatically found the pixels carrying the most information, and de-emphasized background pixels. Alternatively, we used a user-defined region of interest and calculated fluorescence from the unweighted mean of pixel values within this region. For calculations of Δ*F/F*, background fluorescence from a cell-free region was subtracted from the baseline fluorescence of the cell.

### Sub-frame interpolation of AP timing (SNAPT)

Dendritic propagation spike width and delay were calculated using the SNAPT algorithm as described previously.^1^ Briefly, spikes were identified from the whole-cell fluorescence trace. Spike timing was used to construct a spike-triggered average movie. The mean spike waveform was then used as a template and fit to the spike waveform at each pixel, using as fitting parameters: vertical offset, amplitude, time-shift, and width (uniform dilation in time). The time-shift and width parameters are displayed in Fig. 2k.

### Extracting fluorescence from multicellular movies in behaving animals

Movies were first corrected for motion using the NoRMCorre algorithm.^21^ Movies were then corrected for photobleaching by dividing the movie by an exponential fit of the mean fluorescence. Activity-based image segmentation was performed on the spiking component of the signal. To remove subthreshold signals for segmentation purposes, movies were high-pass filtered in time with a 50 Hz high-pass filter. Movies were then segmented semi-automatically using principal components analysis followed by time-domain independent components analysis (PCA/ICA).^22^ ICA produces an arbitrary number of candidate cells. We set the maximum number of cells to 15 and then further eliminated traces that did not correspond to cells by manually inspecting the traces and corresponding spatial maps. In some cases we divided the movie into sub-movies based on patterns of illumination from the DMD masks (Fig. 3d), and performed the PCA/ICA analysis separately in each sub-movie. The spatial masks from PCA/ICA were then applied to the movies without high-pass filtering to extract fluorescence traces that included subthreshold dynamics. Some morphologically distinct cells did not spike during the recording session, consistent with previous whole cell patch clamp recordings in the CA1^23^,^24^, and were excluded from the analysis. Due to cell-to-cell variations in DC background level, the offset of fluorescence traces was not judged to be meaningful. Traces were mean-subtracted and then divided by their standard deviation.

### Spike detection, correlations and spike-triggered averaging

A simple threshold-and-maximum procedure was applied for spike detection. Fluorescence traces were first high-pass filtered, and initial threshold was set at 3 times the noise level. This threshold was then manually adjusted if needed. All analyses of brain state-dependent activity (Fig. 4) were performed on segments of the recordings comprising the last 9 s of the first rest period (quiet) and the first 9 s of the first walk period.

Fluorescence signals were first converted to ΔF/F_0_ where F_0_ was the mean fluorescence in the epoch. The data were mean-subtracted in 1 s windows to correct for slow drifts in baseline fluorescence. To estimate the statistical errors in the cross correlations, we calculated the cross-correlations between randomly selected cells from distinct fields of view.

Spike triggered averages of the fluorescence (Figs. 4g, S8d) were calculated for each pair of simultaneously recorded cells, normalized to the range 0–1, and then averaged across all cell pairs. Spike triggered histograms of spike timing (Fig. 4h) were constructed for each spike with 10 ms bins. These histograms were then summed over all spikes and all cells and divided by the total number of spikes to yield a percentage of spikes in each time bin.

